# The “Gini index” in genetics: measuring genetic architecture complexity of quantitative traits

**DOI:** 10.1101/018713

**Authors:** Xia Shen

## Abstract

Genetic architecture is a general terminology used and discussed very often in complex traits genetics. It is related to the number of functional loci involved in explaining variation of a complex trait and the distribution of genetic effects across these loci. Understanding the complexity level of the genetic architecture of complex traits is essential for evaluating the potential power of mapping functional loci and prediction of complex traits. However, there has been no quantitative measurement of the genetic architecture complexity, which makes it difficult to link results from genetic data analysis to such terminology. Inspired by the “Gini index” for measuring income distribution in economics, I develop a genetic architecture score (“GA score”) to measure genetic architecture complexity. Simulations indicate that the GA score is an effective measurement of the complexity level of complex traits genetic architecture.

## Introduction

*Genetic architecture* has been widely adopted as a general terminology that defines how a complex trait is regulated terminology that defines how a complex trait is regulated by the genome. Interestingly, although the concept is easily defined, we know little about the genetic architecture of any complex trait. Uncovering the underlying genetic architecture, step by step, therefore becomes the main goal of quantitative genetics research.

Although the genetic architecture of each measured complex trait is unknown, it is always needed to further understand the estimates derived from the data (e.g. heritability) based on potential genetic architecture complexity. For instance, two complex traits that have the same estimated narrow-sense heritability (*h*^2^) may have very different genetic architecture, as the number of genetic variants contribute to such an *h*^2^ value may differ substantially. In general, the more polygenic a trait is, the more difficult it is to map functional loci using genomewide association studies (GWAS) [1].

Thus, it is particularly useful to develop a general statistic that measures the complexity of a complex trait, given its phenotypic measurements and genotypes in a population. The genetic architecture complexity describes how polygenic a trait is, namely, how evenly the genetic variance is distributed across all the genotyped variants. Given a certain null hypothesis, mimicking the well-known economic concept “Gini index” [2] that measures the distribution of income in a population, developed here is a score that measures the complexity level of genetic architecture as the distribution of genetic variance in a genome. Simulations indicate that the developed score is an effective measurement.

## Methods

### Statistical modeling

Prior to the development of the GA score, the genetic architecture complexity of a quantitative trait needs to be defined using the phenotype and genome-wide genotype data. Consider the linear whole-genome regression model

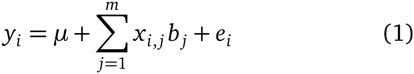

where *y*_*i*_ is the phenotypic value of individual *i*, *x*_*i*,*j*_ its genotypic value of SNP *j* coded as −2 *f* _*j*_, 1 − 2 *f* _*j*_, 2 − 2 *f*_*j*_ (*f*_*j*_ is the allele frequency of SNP *j*), *b*_*j*_ the SNP effect, and *e*_*i*_ the residual. The weights of different SNP effects can be modeled as

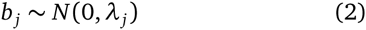

Such a hierarchical model defined by eq. (1) and (2) is actually a double hierarchical generalized linear model [3] for high-throughput genetic markers [4]. The main idea of most current whole-genome regression methods is to optimize the estimation of the weights *λ*_*j*_’s, as the markers are ought to be re-weighted differently for different traits due to different genetic architectures.

### Genetic architecture score

Let us define the genetic architecture complexity level as the *weight distribution* of genome-wide SNP effects, so that

- The highest complexity: *λ*_*j*_ is uniformly distributed across all the markers;
- The lowest complexity: *λ*_*j*_ *>* 0 and *λ*_*j* (*j′* ≠*j′*)_ = 0, i.e. only one marker is predictive of the trait.

Based on a null hypothesis of the highest complexity, we have

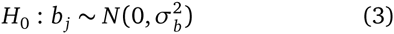

Eq. (1) becomes a ridge regression, a.k.a. SNP-BLUP model [5, 6]. Under the null hypothesis, every marker is predictive of the trait, so that for SNP *j*, after training the SNP-BLUP model in a training set, the variance explained by SNP *j* in a test set is proportional to its expected value. Namely, we have

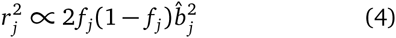

where 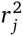 is the variance captured by SNP *j* in the test set, *f* _*j*_ the allele frequency of SNP *j* in the test set, and 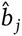 the estimated SNP effect in the training set.

Due to shrinkage estimates in high-dimensional genotype data, one cannot simply add up 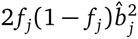 from SNPBLUP for multiple SNPs to obtain the cumulative expected variance explained by a group of SNPs. Nevertheless, it is possible to mimic the Gini index definition: Ranking the SNPs, from the lowest to the largest expected variance explained, one can calculate the observed cumulative variance explained by the first *k* SNPs is

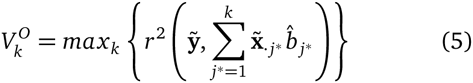

where *j\** represents the rank, 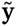 and 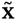 are the phenotype and genotype vectors in the test sample, and *r* stands for correlation coefficient. Under the null, 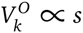 where *s* is the number of SNPs included as predictors. A deviation from the null hypothesis will result in a function *V*^O^ = *g*(*s*) deviating from the straight line *V*^O^ = *cs* (Figure 1), where *c* = 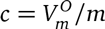. Therefore, the area between the curve *V*^O^ = *g*(*s*) and the line *V*^O^ = *cs* measures the genetic architecture complexity.

**Figure 1.**
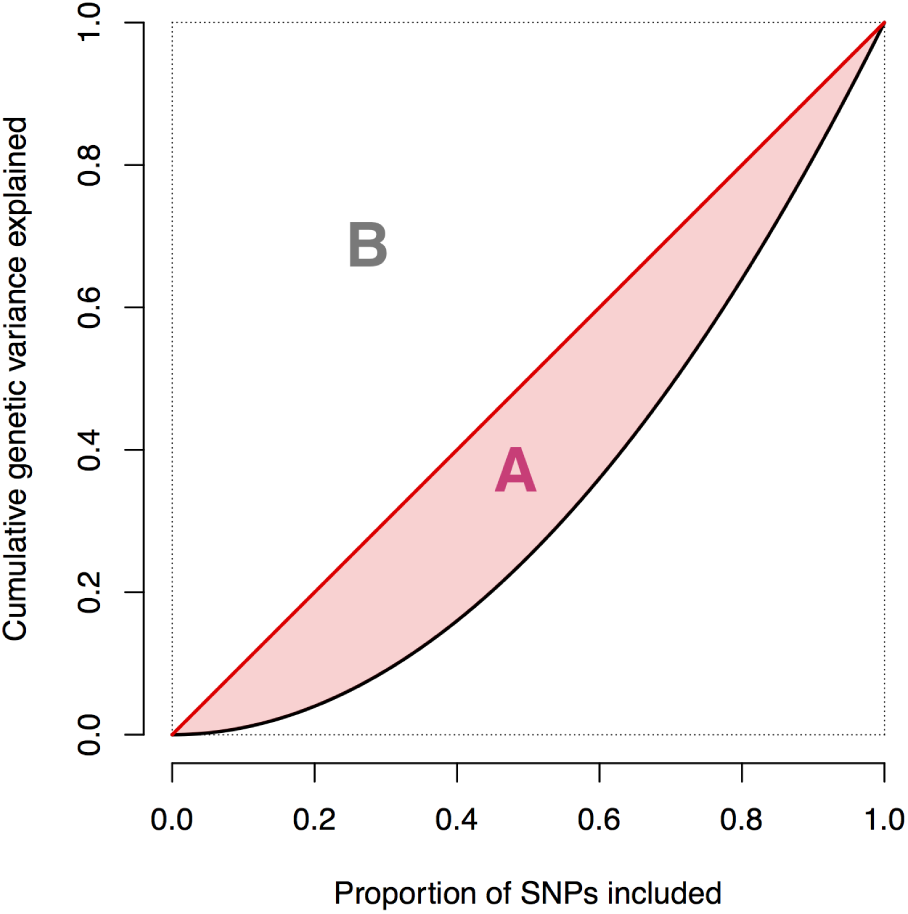
Illustration of the GA score definition. The black curve is plotted with 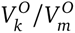 against 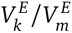. The GA score is defined as the ratio (*B A*)*/B*, where the area *A* varies from zero (null: highest complexity) to *B* = 0.5 (lowest complexity: “monogenic”).

A genetic architecture (GA) score can be defined as an area ratio of 1 *− A/B* illustrated in Figure 1, i.e.

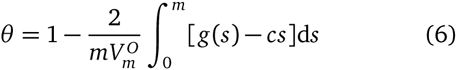

which ranges from 0 (lowest complexity: “monogenic”) to 1 (null: highest complexity).

## Results

Three phenotypes were simulated based on the simulated genotypes announced by the Genetics Society of America (GSA) [7], for both the training set (*n* = 2 000) and validation samples (simulated young generations, *n* = 1 500). All the three phenotypes had exactly the same simulated narrow-sense heritability (*h*^2^ = 0.5) but different numbers of causal variants (50, 5 000, 50 000, respectively). The whole genome contains 57 458 genotyped markers in total.

A SNP-BLUP model was fitted for each phenotype using the *bigRR* package [6] in the training population. The estimated SNP effects were passed onto the validation set to compute the GA score *θ*. The estimated GA scores were 0.42, 0.85 and 0.93 for the 50-, 5 000- and 50 000-causal-variants architectures, respectively. So the estimated GA score grows, although not linearly, with the number of causal variants.

## Discussion

The GA score developed here measures the genetic architecture complexity in terms of the distribution of genetic variance over the genome. Nevertheless, the estimated scores are more useful when comparing multiple phenotypes. An interesting additional study is to correlate a score as such to the sample size required in GWAS meta-analysis of a certain trait, so that one can predict the chance of new discoveries in GWAS prior to data collection.

**Figure 2.**
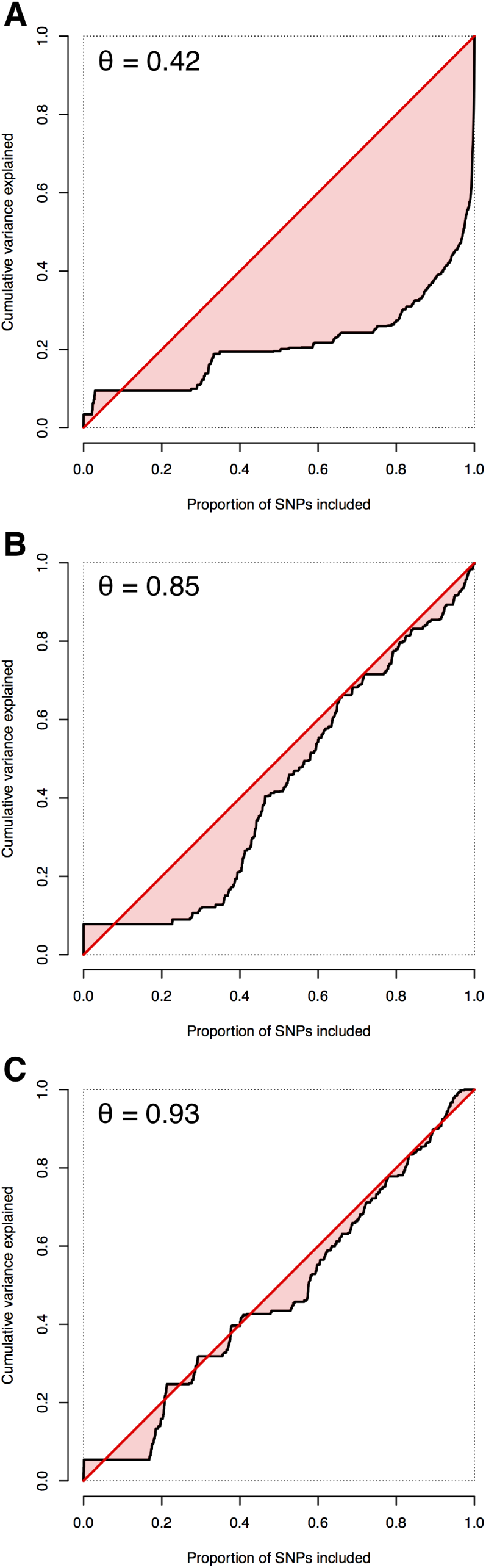
GA score applied on three different scenarios of the GSA simulated data. A narrow-sense heritability of 0.50 was simulated in all scenarios. 50 (A), 5 000 (B), and 50 000 (C) causal markers were simulated, respectively.

### Author contributions

X.S. initiated the study, developed the method, analyzed the data, wrote and approved the final version of the manuscript.

### Grant information

This work is funded by a Swedish Research Council grant (2014-371) to X.S..

## Acknowledgements

The author thanks Chris S. Haley and Ziqing Weng for helpful input.

## References

[1] Peter M. Visscher, Matthew A. Brown, Mark I. McCarthy, and Jian Yang. Five years of gwas discovery. American Journal of Human Genetics, 90:7–24, 2012.

[2] C. Gini. On the measure of concentration with special reference to income and statistics. Colorado College Publication, General Series, (208):73–79, 1936.

[3] Y. Lee and J. A. Nelder. Double hierarchical generalized linear models (with discussion). Applied Statistics, 55: 139–185, 2006.

[4] X. Shen, L. Rönnegård, and O. Carlborg. Hierarchical likelihood opens a new way of estimating genetic values using genome-wide dense marker maps. BMC Proceedings, 5 (Suppl3)(S14), 2011.

[5] HE Meuwissen, BJ Hayes, and ME Goddard. Prediction of total genetic value using genome-wide dense marker maps. Genetics, 157(4):1819–1829, APR 2001.

[6] X Shen, M Alam, F Fikse, and L Rönnegård. A novel generalized ridge regression method for quantitative genetics. Genetics, 193(4):1255–1268, April 2013.

[7] J. M. Hickey and G. Gorjanc. Simulated data for genomic selection and genome-wide association studies using a combination of coalescent and gene drop methods. G3: Genes|Genomes|Genetics, 2(6), 2012.

